# Cross-talk between tissues is critical for intergenerational acclimation to environmental change

**DOI:** 10.1101/2023.11.15.567297

**Authors:** Sneha Suresh, Megan J Welch, Philip L Munday, Timothy Ravasi, Celia Schunter

## Abstract

An organism’s reaction to environmental changes is mediated by coordinated responses of multiple tissues. Additionally, parental priming may increase offsprings’ acclimation potential to changing environmental conditions. As the effects of human-induced climate change, such as ocean acidification (OA), continue to intensify it is critical to assess the acclimation potential of species at the whole organismal scale. For this we need to understand the cross-talk between tissues in regulating and responding to pH changes. Here by using a multi-tissue approach we determine the influence of 1) variation in parental behavioural tolerance and 2) parental environment, on molecular responses of their offspring in a coral reef fish. The gills and liver showed the highest transcriptional response to OA conditions in juvenile fish regardless of the parental environment, while the brain and liver showed the greatest signal of intergenerational acclimation. Key functional pathways that were altered in the brain upon within-generational CO_2_ exposure were restored to control levels when parents were exposed to OA conditions. Furthermore, the expression of a new complement of genes involved in key functions was altered in the offspring only when parents were previously exposed to OA conditions. Therefore, previous parental conditioning to OA can reprogram tissue transcriptomic profiles of the offspring enabling them to better cope in an environment with elevated CO_2_ levels. Overall, our results reveal tissue-specific transcriptional changes underlying intergenerational plastic responses to elevated CO_2_ exposure and highlight the integration of these changes in promoting organismal acclimation to OA.

## Introduction

With the global climate continuously shifting to more extreme conditions organisms need to acclimate and/or adapt to the changing environments in order to survive. The oceans are becoming increasingly acidified as they absorb a major portion of anthropogenic CO_2_ emissions (Pörtner et al., 2022) leading to ocean acidification (OA) which is reported to negatively impact the physiology and behavior of various marine organisms including fish (Heuer & Grosell, 2014; Strader et al., 2020). However, increasing evidence suggests that multi-generational exposure to elevated CO_2_ conditions could influence the adaptive capacity of future generations to OA conditions (Nagelkerken et al., 2023). In fact, several studies have reported transgenerational acclimation in a number of fish species as well as some invertebrates to OA (Strader et al., 2020). Specifically, transgenerational exposure to elevated CO_2_ conditions has been shown to facilitate acclimation of metabolism, growth, survival, neuronal plasticity and behavior in independent studies (Allan et al., 2014; Miller et al., 2012; Monroe et al., 2021; Munday, 2014; Schade et al., 2014; Schunter et al., 2016, 2018; Stiasny et al., 2018) however, we are still learning about the underlying molecular mechanisms of such acclimation processes.

Additionally, variation both within and across species in the biological responses to OA also exists due to differences in their evolutionary and environmental history. Studies examining the effect of elevated CO_2_ on metabolism, growth, development, and reproduction in fish show variable results with some species being more affected than others (Heuer & Grosell, 2014). Variation in sensitivity to elevated CO_2_ within a population could be crucial in long-term adaptation through selection of more tolerant individuals. Indeed, individual variation in behavioural tolerance to elevated CO_2_ exposure has been reported to be heritable and hence could facilitate rapid selection of tolerant genotypes in the population (Lehmann et al., 2022; Welch & Munday, 2017). Such selection for CO_2_ tolerance has been shown to occur in nature, which could result in populations consisting of individuals with greater behavioural tolerance to elevated CO_2_ (Munday et al., 2013). Furthermore, inter-individual variation in sensitivity to ocean acidification could have an epigenetic basis (Ryu et al., 2018; Turner, 2009) and in fact several studies have reported the expression levels of genes involved in epigenetic processes to be altered upon exposure to elevated CO_2_ conditions (Huang et al., 2019; Schunter et al., 2018). Transfer of epigenetic factors from parents to offspring (epigenetic inheritance) could be one of the potential mechanisms of inter-and trans-generational acclimation and eventual adaptation to OA.

Adaptive processes to environmental changes at the organismal level require integrated activity of various tissues, with each tissue undergoing changes in its transcriptional landscape resulting in the overall response of the organism. However, to date, research has mainly focused on individual tissue functional changes in response to OA with less emphasis on how these changes integrate to create a whole-body response. Several studies have examined the effects of OA on brain and neurosensory systems since the discovery of impaired behavioural responses in various fish species in elevated CO_2_ conditions. The altered behavioural responses have been linked to changes in the functioning of the GABAergic signaling pathway (Schunter et al., 2019) and the circadian rhythm in the brain of fish exposed to elevated CO_2_ (Lee et al., 2021; Schunter et al., 2016; Williams et al., 2019). Previous studies have also focused on the effects of OA on the gill transcriptome due to it being the primary organ involved in acid-base regulation, immune defense, and stress response, and hence plays a vital role in maintaining cellular homeostasis under conditions of CO_2_ stress (De Souza et al., 2014; Deigweiher et al., 2008, 2010). These processes are energetically expensive and indeed changes in the aerobic metabolic scope (Crespel et al., 2019; Gräns et al., 2014; Pimentel et al., 2014; Rummer et al., 2013) and expression levels of key metabolic genes (Frommel et al., 2020) have been reported in fish exposed to elevated CO_2_. Therefore, exposure to elevated CO_2_ affects various aspects of fish physiology such as metabolism, cellular redox status, ion transport and acid-base homeostasis, neurological functioning and behavior thereby exerting a whole-body functional reprogramming (Grosell et al. 2019). Therefore, a systematic transcriptomic analysis is needed to determine how the biological processes associated with each tissue integrate together within the whole organism to drive adaptive responses to elevated CO_2_ environments.

In this study we conducted an intergenerational CO_2_ exposure experiment and performed systematic analysis of gene expression changes in response to elevated CO_2_ across three tissues, the brain, the gills, and the liver, in the spiny damselfish *Acanthochromis polyacanthus*. While this species can be sensitive to increases in water temperature and CO_2_ levels, they have the potential to acclimate to the changing environmental conditions across multiple generations (Donelson et al., 2012; Schunter et al., 2016, 2018; Veilleux et al., 2015). *A. polyacanthus* has been used as a model to study the impacts of climate change, and to investigate the molecular basis of intergenerational plasticity to environmental changes, due to its advantageous life-history traits for laboratory studies (Robertson, 1973), however past studies have only focused on single tissues (Ryu et al., 2018; Schunter et al., 2016, 2018). Here, by using a multi-tissue transcriptomic approach we aim to determine how dynamic cross-talk between tissues maintains whole-body homeostasis under future ocean acidification conditions. Additionally, we also assess how the acclimatory response of offspring mediated by transcriptional reprogramming across multiple tissues is influenced by variation in parental sensitivity to elevated CO_2_ and parental environment. Through systemic characterization of the effects of OA we aim to identify how the adaptive processes within each tissue integrate to drive intergenerational acclimation to OA at the organismal level.

## Methods

### Sample collection, behavioural testing, and experimental design

Adult *Acanthochromis polyacanthus* were collected from the wild on the Great Barrier Reef, Australia (18°38’24.3’S, 146°29’31.8’E) and exposed to elevated CO_2_ (754 ± 92 µatm) for seven days following which their behavioural sensitivity to conspecific chemical alarm cues (CAC) was tested using a two-chamber flume as described previously (Schunter et al., 2016). Briefly, the fish were classified as being behaviorally sensitive or tolerant to elevated CO_2_ based on the amount of time spent in water containing the CAC. Sensitive individuals spent ≥ 70% time in CAC whereas tolerant individuals spent ≤ 30% time in CAC. Individuals of similar size displaying the same behavioural phenotype (sensitive or tolerant) were then grouped into breeding pairs and held in either control (414 ± 46 µatm) or elevated CO_2_ conditions (754 ± 92 µatm) for three months prior to the breeding season. Offspring clutches from each breeding pair were placed into three different experimental treatments resulting in three combinations of parent-offspring conditions for each parental phenotype: (1) Control treatment – Parents and offspring held at control condition (414 ± 46 µatm); (2) Developmental treatment – Parents held at control condition and offspring exposed to elevated CO_2_ (754 ± 92 µatm) immediately after hatching; and (3) Intergenerational treatment – Parents and offspring exposed to elevated CO_2_ (754 ± 92 µatm). The offspring were held in their respective conditions until they were five months old after which nine fish from each parental phenotype, from each treatment condition (N = 27 from each parental phenotype; N = 54 total fish sampled) were euthanized and the brain, gills and liver were dissected, snap frozen in liquid nitrogen and stored at −80 °C until further processing (Supplementary Figure S1).

### RNA extraction, sequencing, and gene expression analyses

Total RNA was extracted from the fish brains, livers and gills using the AllPrep DNA/RNA Mini kit from Qiagen following the manufacturer’s instructions. RNA quality was determined using nanodrop and Agilent Bioanalyzer and samples having an RNA integrity value (RIN) ≥ 8 were sequenced using Illumina HiSeq 2500 to get paired-end reads of 100 bp at Macrogen Inc., South Korea. A total of 1,614.25 ± 3.05, 2,367.03 ± 5.19, and 2,227.23 ± 6.37 million raw paired-end reads were obtained from the 162 sequenced libraries from brain, gills and liver, respectively which included nine control, nine developmental and nine intergenerational samples for each parental phenotype for each tissue (Supplementary Table S1). The quality of the raw reads were examined using FastQC (Andrews, 2010) v0.11.8 and adapters and low quality sequences were trimmed using Trimmomatic (Bolger et al., 2014) v0.39 (ILLUMINACLIP: adapters.fa:2:30:15:8:true; SLIDINGWINDOW:4:20; MINLEN:32).

Only those sequences ≥ 32 bp in length with both the forward and reverse reads retained after trimming were used for further analysis. Potential contaminant sequences were identified using kraken (Wood & Salzberg, 2014) v2.0.8-beta, with a confidence score of 0.3, using the bacteria, fungi and virus RefSeq genomic libraries as reference and removed from further analyses. A total of 1,510.51 ± 2.62, 2,254.16 ± 4.95, and 2,116.25 ± 6.04 million high-quality sequences were retained after the filtering process (Supplementary Table S1). These sequences were mapped to the *Acanthochromis polyacanthus* reference genome (unpublished) using HISAT2 (Kim et al., 2019) v2.1.0. On average, 84 ± 1.83%, 91.22 ± 0.66%, and 93.33 ± 0.81% reads mapped to the reference genome from the brain, gills, and liver respectively (Supplementary Table S1). Raw read counts per gene were obtained using featureCounts (Liao et al., 2014) v2.0.0 (parameters: -B -J -M --fraction), assigning fractional counts to multi-mapped reads. Exploring the gene expression patterns across the whole dataset (162 samples) using principal component analysis (PCA) revealed a clear clustering of samples by tissues indicating that tissues vary greatly in their gene expression patterns (Supplementary Figure S2). Subsequent analysis of differences in gene expression levels was therefore carried out separately for each tissue using the DESeq2 (Love et al., 2014) v1.32.0 package in R (R Core Team, 2021) v4.2.1.

Principal component analysis (PCA) using the regularized log transformed (rlog) counts was done in R v4.2.1 to detect and remove outlier samples. A likelihood ratio test (LRT) using a model comparison approach was then used to determine the effect of family line in driving the gene expression patterns and to determine the best design formula for the final DE analysis. First, significant differences in gene expression were measured by comparing a model including treatment and family line against a reduced model without the family line factor separately for each tissue. For a total of 924, 910, and 923 genes in the brain, gills, and liver respectively, the model including family line better explained the observed differences in gene expression compared to the reduced model excluding this factor (FDR corrected p-value < 0.05; Supplementary Table S2). Pair-wise comparisons between the control, developmental and intergenerational treatment was then caried out in DESeq2 (accounting for the family effect, using Wald test) separately for each parental phenotype for each tissue to determine the effect of parental environment and parental tolerance to CO_2_ on the molecular responses of the offspring to elevated CO_2_. For each pair-wise comparison, the genes were considered to be significantly differentially expressed (DE) if the False Discovery Rate (FDR) adjusted p-value was less than 0.05, the absolute log 2-fold change in expression was greater than 0.3 and baseMean was greater than 10. Functional enrichment analysis of the significant DE genes was carried out in OmicsBox (https://www.biobam.com/omicsbox) v1.4.11 using Fisher’s Exact Test (FDR corrected p-value < 0.05) with the option of reducing to most specific GO terms to reduce redundancy. The genes associated with the enriched GO terms were further categorized into broader functional groups based on their functional description from the UniProt knowledgebase (UniProtKB; https://www.uniprot.org/). All figures are made using ggplot in R v4.2.1.

## Results

### Molecular processes affected by all elevated CO_2_ treatments

To understand a general effects of elevated CO_2_ exposure regardless of time of exposure to elevated CO_2_, we identified genes that were commonly differentially expressed (DE) in both the developmental and intergenerational treatments compared to control (Figure 1(a)(i)), which are considered the general “CO_2_ response genes”. Irrespective of the parental environment, the liver showed the greatest transcriptional response to elevated CO_2_ exposure in offspring of both sensitive and tolerant parental behavioral phenotypes, with 63 and 986 differentially expressed (DE) genes, respectively (Figure 1(b); Supplementary Table S4). In the offspring of tolerant parental phenotype, the gills followed with 883 DE genes, while the brain had 44 DE genes (Figure 1(b)). Conversely, in the offspring of sensitive parental phenotype, the brain displayed a higher number of DE genes (37) compared to the gills (9). Overall, offspring of parents with tolerant behavioural phenotype showed increased capacity for gene expression regulation across all three tissues in response to elevated CO_2_ exposure. However, the CO_2_ response genes showed high tissue specificity with no genes being commonly DE across the three tissues in the sensitive parental phenotype and only 3 and 81 genes being shared between the brain and gills, and liver and gills respectively in the tolerant parental phenotype (Supplementary Figure S3(a)).

**Figure 1:**
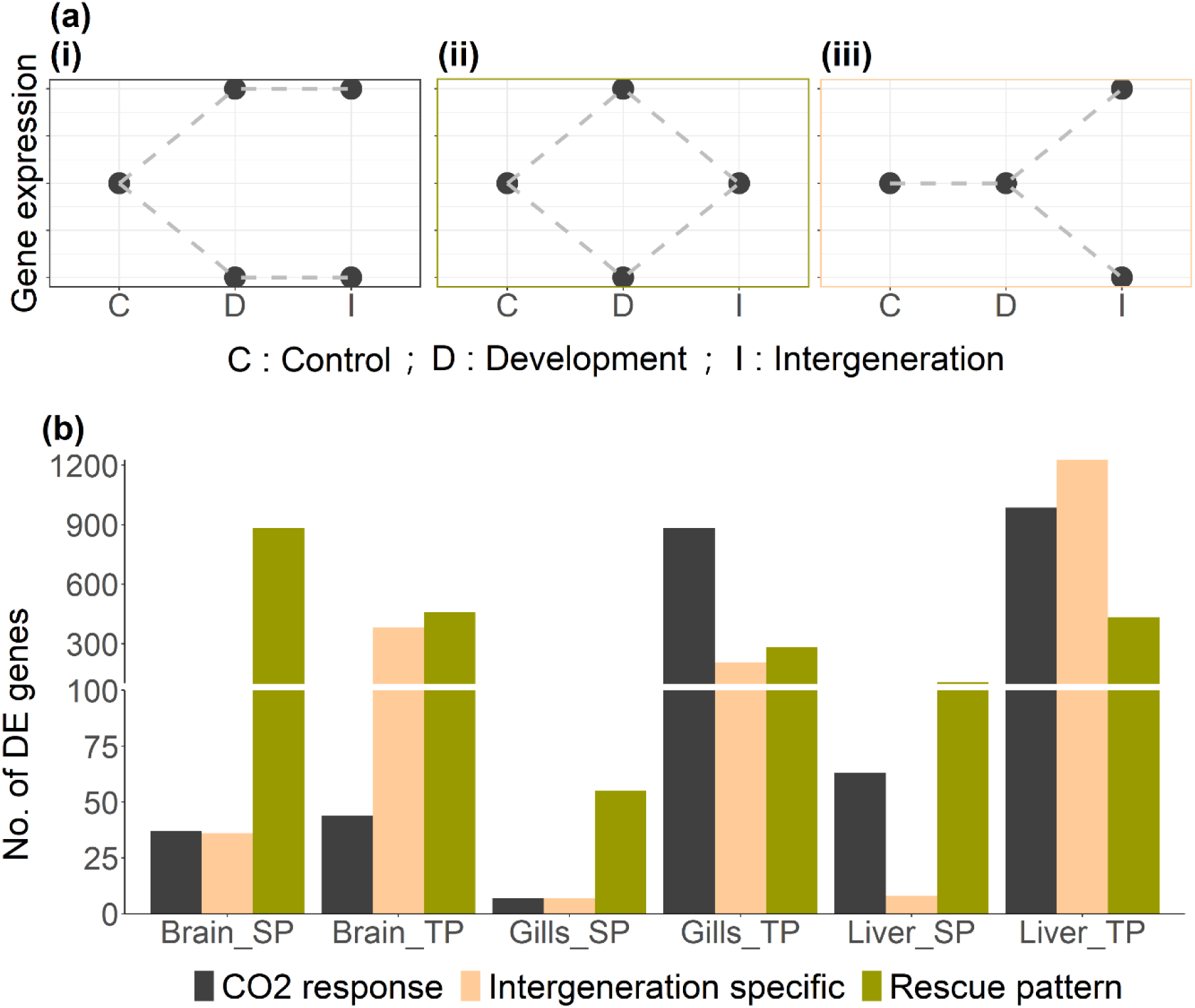
(a) Schematic graph representing the expression profile of (i) CO_2_-response genes, (ii) genes showing a rescue pattern, and (iii) intergeneration-specific genes. (b) Number of differentially expressed genes across the three treatments in all the three tissues. SP indicates samples with a sensitive parental phenotype and TP indicates samples with a tolerant parental phenotype for each of the respective tissues. Note scale break in y-axis at 100 DE genes.

Functional enrichment analysis of the CO_2_ response genes in offspring with sensitive parental phenotype revealed translation and amino-acid synthesis to be important in the brain, while no significantly enriched functions were found in the gills and liver (Figure 2; Supplementary Table S5). For the offspring of tolerant parental phenotype, biosynthetic processes was commonly enriched in the brain and gills (Figure 2). Few other functional pathways were commonly enriched between the gills and liver, including metabolism, translation, protein transport, immune and stress response (Figure 2; Supplementary Table S5). Interestingly, while genes associated with metabolic pathways such as glycolysis, TCA cycle and mitochondrial electron transport chain were commonly differentially regulated in both the gills and liver, pentose phosphate pathway gene showed upregulation exclusively in the gills. Additionally, immune response and translation showed tissue-specific regulation with downregulation in the liver but upregulation in the gills (Supplementary Table S5). Furthermore, RNA processing and primary active transmembrane transport were enriched only in the liver (Supplementary Table S5). This suggests intricate and tissue-specific regulation of various functions in response to elevated CO_2_ exposure.

**Figure 2:**
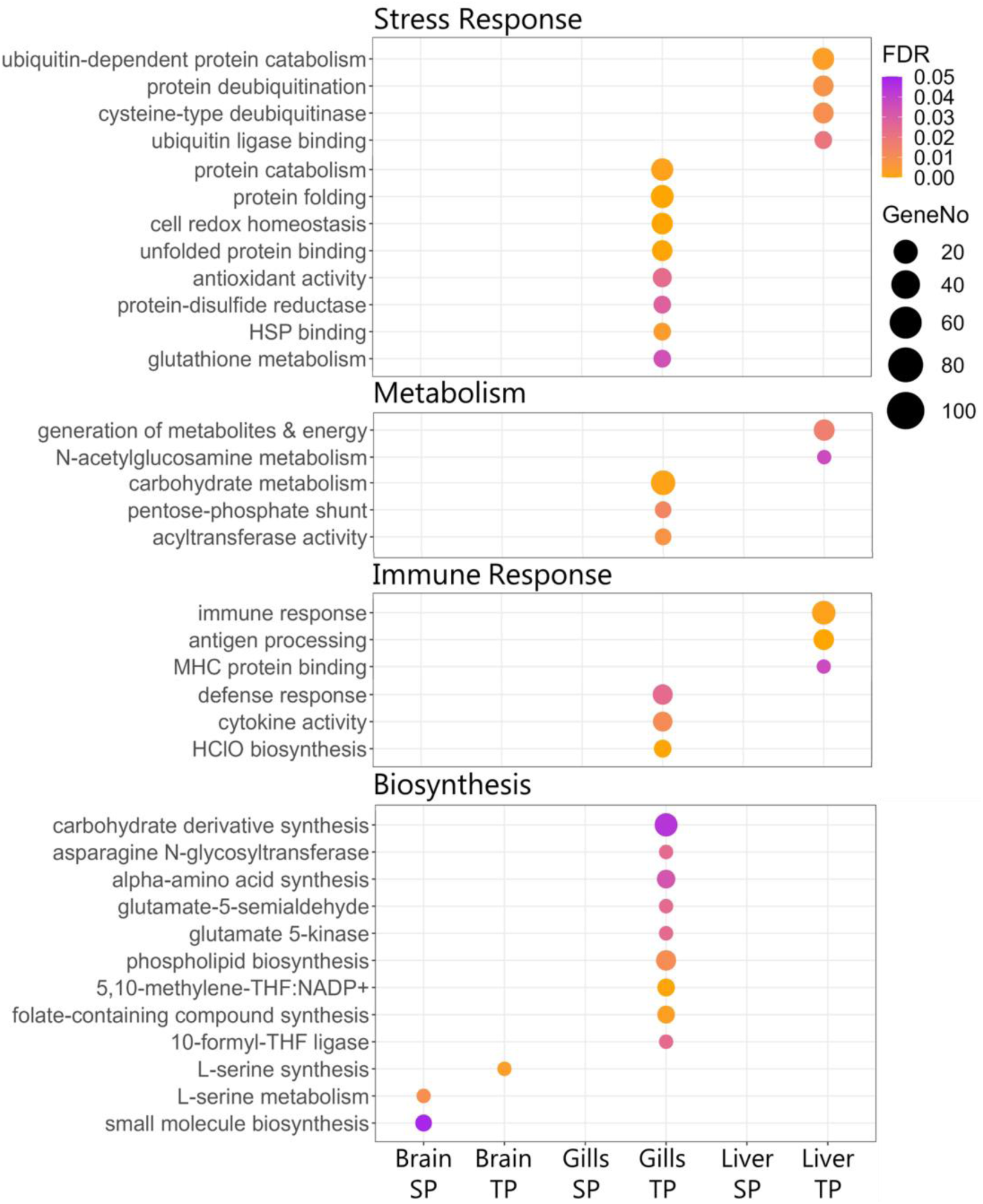
Functions that are significantly enriched (FDR <0.05) among the DE genes involved in overall CO_2_ response. SP indicates offspring of behaviorally sensitive parents and TP indicates offspring of behaviorally tolerant parents. The color of the circles represents the FDR corrected p-value and size of the circles represents the number of genes associated with the respective function.

## Parental exposure to elevated CO_2_ “rescues” developmental effects

In the developmental treatment (compared to control and intergeneration) a total of 883, 55, and 108 genes that were DE in offspring with sensitive parental phenotype and 458, 284, and 434 genes that were DE in offspring with tolerant parental phenotype in the brain, gills, and liver respectively, returned to control levels when parents were previously exposed to OA conditions suggesting cross-generation plasticity resulting in “rescue” of gene expression levels (Figure 1(a)(ii); Supplementary Table S6). Parental conditioning had the largest effect on brain gene expression followed by the liver (Figure 1(b)). Similar to the CO_2_-response genes, there were very few genes commonly DE across tissues (Supplementary Figure S3(b)).

Genes exhibiting a rescue pattern in the brain of offspring of behaviorally sensitive parents showed the highest degree of specificity in functional regulation with pathways involved in cytoskeleton, apoptosis, synaptic signaling, sodium and calcium channel activity, transcription regulation, cellular transport, and small GTPase signaling being exclusively enriched only in this tissue (Figure 3; Supplementary Table S7). Gills were found to play a key role in regulating ion transport in offspring of both behaviorally sensitive and tolerant parents. Specifically, the expression of genes encoding sodium and potassium channel transporters were altered in the gills upon developmental exposure to elevated CO_2_ levels but returned to control levels in intergenerationally exposed fish (Figure 3; Supplementary Table S6, S7). Additionally, purine ribonucleoside salvage pathway and DNA replication were identified as key functions among the “rescue” genes in the offspring of parents with sensitive phenotype in the gills and liver respectively (Figure 3; Supplementary Table S7). This suggests that the molecular response of the offspring is influenced by the parental environment with specific functional pathways with parental conditioning to elevated CO_2_ resulting in selective regulation of specific functions in each tissue.

**Figure 3:**
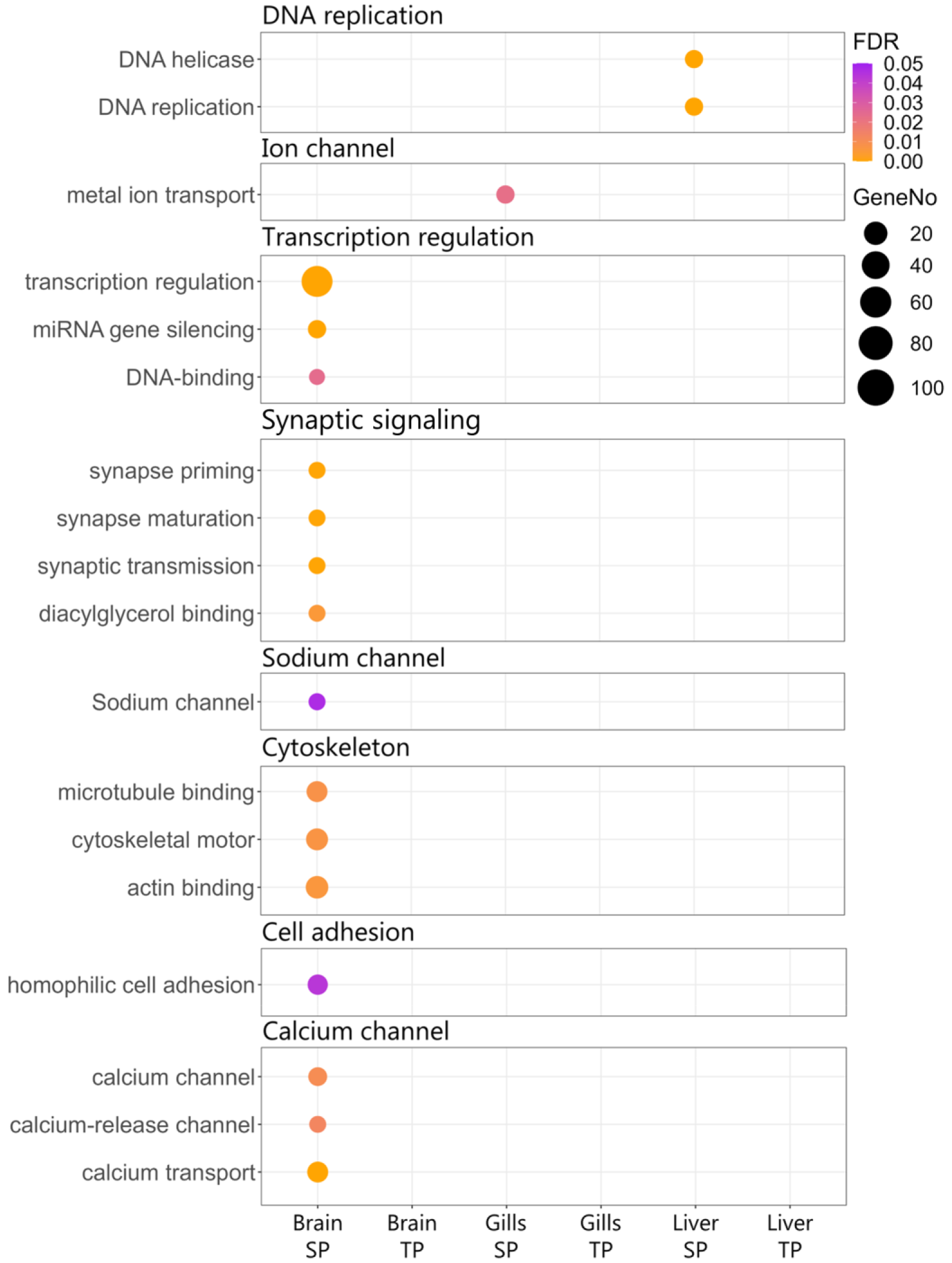
Functions that are significantly enriched (FDR <0.05) among the DE genes showing a rescue pattern. SP indicates offspring of behaviorally sensitive parents and TP indicates offspring of behaviorally tolerant parents. The color of the circles represents the FDR corrected p-value and size of the circles represents the number of genes associated with the respective function.

### Intergenerational specific response to elevated CO_2_

We found a large transcriptional response to elevated CO_2_ that was only seen in the intergenerationally exposed fish and not in fish with only developmental (within generation) exposure to elevated CO_2_. This indicates plasticity of the offspring transcriptome due to parental conditioning to elevated CO_2_ and was especially marked in offspring of tolerant parents (Figure 1(b)). These are genes that were only differentially expressed in the intergenerational treatment (compared to control and development) but were at control levels in the developmental treatment (Figure 1(a)(iii)). Specifically, 383, 207, and 1,226 genes were DE in the brain, gills, and liver respectively in offspring with tolerant parents, while offspring with sensitive parents had 36, 7, and 8 DE genes in the brain, gills, and liver respectively that were specific to the intergenerational treatment (Supplementary Table S8). Offspring of both tolerant and sensitive parents showed high tissue specificity in the intergenerational specific transcriptional signature to elevated CO_2_ with only one and five genes being commonly DE across all three tissues in the sensitive and tolerant parental phenotypes respectively (Supplementary Figure S3(c)).

The intergeneration specific response was substantially higher in offspring of behaviourally tolerant parents with various functions showing tissue-specific regulation. Stress response and metabolic pathways were commonly differentially regulated in the brain and liver (Figure 4), but the liver showed a much larger number of DE genes (∼400) associated with metabolism compared to the brain (12). Additionally, signal transduction pathway was enriched only in the brain while several functions such as biosynthetic pathways, endopeptidase activity, lipid binding/ transport, transcriptional regulation, proteolysis, and spindle organization were exclusive to the liver (Figure 4, Supplementary Table S9). Interestingly, molecular signatures indicating pH regulation by bicarbonate retention were identified only in the intergenerationally treated fish with tolerant parental phenotype, indicated by the downregulation of SLC4A1, CFTR, SLC12A2, and SLC9A3 in the gills and upregulation of SLC4A4 in the brain (Supplementary Table S6). Therefore, offspring of parents with tolerant behavioural phenotype showed increased molecular signatures of intergenerational plasticity and this was especially evident in the liver. This change in transcriptional signature could regulate the above mentioned “rescue” pattern and equip offspring to better cope with OA conditions.

**Figure 4:**
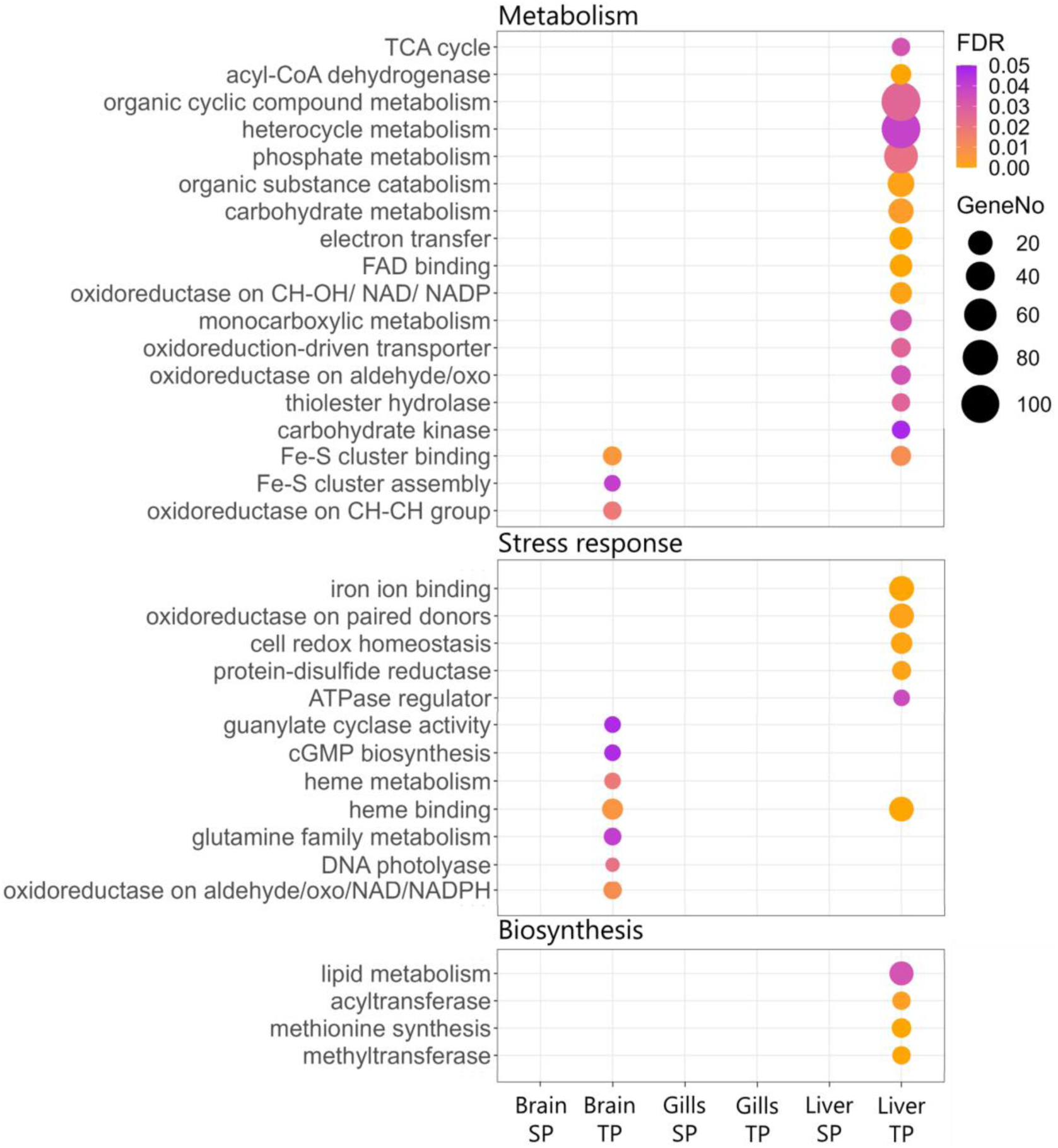
Functions that are significantly enriched (FDR <0.05) among the intergeneration specific DE genes in offspring of behaviorally tolerant parents. The color of the circles represents the FDR corrected p-value and size of the circles represents the number of genes associated with the respective function.

### Parental variability in CO_2_ sensitivity impacts the offspring transcriptome

Parental behavioural phenotype was found to have a substantial influence on the offspring transcriptional response. Across all three tissues, offspring with behaviorally tolerant parents when faced with elevated CO_2_ had larger changes in gene expression levels (log2FC > 5; Figure 5) and a greater number of DE genes involved in the overall CO_2_ response (common in developmental and intergenerational CO_2_ exposure compared to control; Figure 1(b)). There were also very few genes involved in overall CO_2_ response shared between the two parental phenotypes, specifically only six, three, and eleven common DE genes in the brain, gills, and liver respectively (Supplementary Figure S3(a)). When considering genes involved in intergenerational plastic responses, there were more differentially expressed genes in the offspring of tolerant parents compared to those of sensitive parents, except for genes showing a rescue pattern in the brain (Figure 1(b). Additionally, none of the DE genes involved in intergenerational plasticity were shared in the liver tissue between the two parental phenotypes while the brain and gills had a small proportion of common DE genes (specifically, 121 and 11 common DE genes showing a rescue pattern in the brain and gills respectively and only two and one common DE genes in the intergeneration specific response in the brain and gills respectively (Supplementary Figure S3(b, c)).

**Figure 5:**
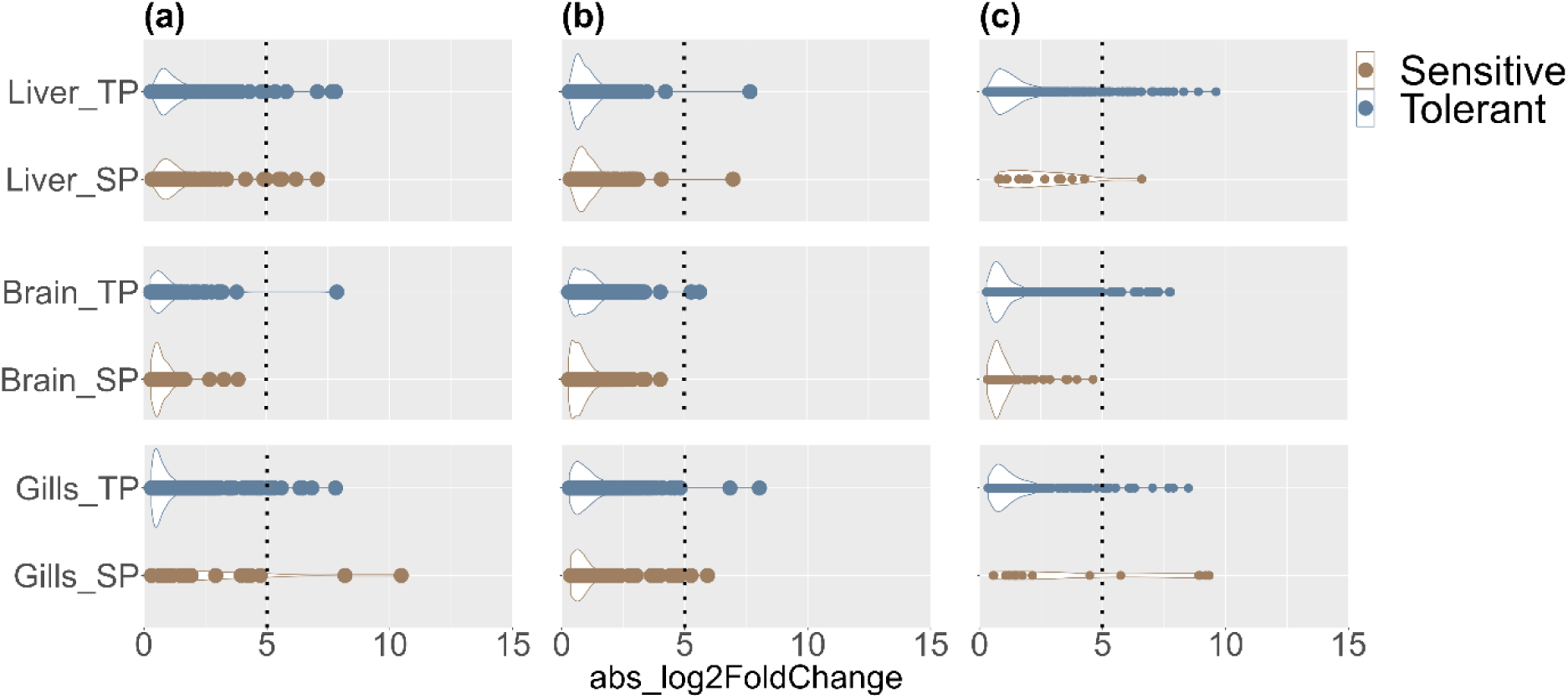
Log2 fold change in expression of genes involved in (a) overall CO_2_ response, (b) rescue pattern, and (c) intergenerational specific response across all three tissues. SP indicates samples with a sensitive parental phenotype and TP indicates samples with a tolerant parental phenotype.

The difference in transcriptional response between the two parental phenotypes was especially pronounced when considering genes showing an intergeneration-specific signature with offspring of tolerant phenotype having more DE genes with a much higher magnitude of gene expression changes (log2FC >5; Figure 5). The liver showed the greatest signal of intergenerational response (Figure 4). Overall, while there were some common transcriptional responses between offspring with sensitive and tolerant parents, offspring of tolerant parents showed a much stronger transcriptional response to elevated CO_2_ in general and also had a stronger signature of intergenerational plasticity.

## Discussion

Changes in the offspring transcriptional landscape in response to elevated CO_2_ exposure exhibited tissue-specific signatures which were largely influenced by previous parental exposure to elevated environmental CO_2_, but also by the parental behavioural phenotype. Specifically, we found that gills are critical in maintaining overall cellular homeostasis in elevated CO_2_ conditions, by regulating pH disturbances and defending against stress-induced cellular damage and potential infections, and the brain and liver had the greatest signal of intergenerational plastic response. In fact, intergenerationally treated fish no longer showed molecular signatures of altered neural signaling in the brain that were seen in the developmental (within generation) treatment. Additionally, differential regulation of a new complement of metabolic genes exclusively in the brain and liver of offspring from parents with prior exposure to elevated CO_2_ implies that parental conditioning enhances the offspring’s capacity to regulate metabolic processes to meet potential shifts in metabolic demands arising from ocean acidification conditions. *A. polyacanthus* is known to have a highly plastic genome enabling it to respond and acclimate to environmental changes (Bernal et al., 2020; Kang et al., 2022) and our results show that this persists across generations potentially enabling this species to rapidly acclimate to the changing ocean environment.

Genes that are always differentially expressed in elevated CO_2_ conditions regardless of the type of exposure are key genes in the general response to ocean acidification (OA). The gills and liver exhibited a higher transcriptional response in all elevated CO_2_ treatments suggesting that these tissues play an important role in the overall response of the fish to elevated CO_2_. In both the developmental and intergenerational CO_2_ treatments, key functional pathways such as cellular metabolism, stress and immune response were differentially regulated in these tissues. Exposure to elevated CO_2_ conditions is known to induce stress and immune response pathways (reviewed in Strader et al., 2020). Interestingly, in our study while stress response genes were upregulated in both gills and liver, genes involved in immune response were upregulated in the gills but downregulated in the liver. This indicates that both the gills and liver work in synergy to mitigate potential cellular damage from oxidative stress and maintain cellular redox balance. However, given that gills are one of the major surface tissues continuously exposed to the external environment, and also serve as a first line of defense against potential infections (Harper & Wolf, 2009; Hu et al., 2023), selective activation of immune responses in this tissue alone could be a preventive measure against opportunistic infections under CO_2_ stress as has been previously observed under elevated CO_2_ conditions (De Souza et al., 2014; Machado et al., 2020). Similar differences in tissue-specific regulation of specific metabolic pathways were observed between the gills and liver in all elevated CO_2_ treatments. Specifically, genes associated with the tricarboxylic acid (TCA) cycle were upregulated in both tissues, but pentose phosphate pathway was upregulated only in the gills. Interestingly, although TCA cycle genes were upregulated in the liver, various genes part of the cytochrome complex, an integral component of the electron transport chain, and genes associated with glycolysis like, Fructose-bisphosphate aldolase B and enolase, were downregulated in the liver. This suggests that there is potentially an overall shift in metabolic priorities and adjustment of ATP production to potentially facilitate redirection of metabolic carbon fluxes to meet cellular demands for various metabolites and cofactors needed for other biological processes such as biosynthetic pathways and the cellular stress response machinery (Gansemer et al., 2020; Rokitta et al., 2012; Walsh & Milligan, 1993). Overall, our results indicate conserved pathways in the gills and liver that are always activated in response to elevated CO_2_ exposure irrespective of previous parental conditions.

Parental exposure to altered environmental conditions can pre-acclimate the offspring transcriptome to these new conditions via intergenerational plasticity. In juvenile *A. polyacanthus*, developmental exposure to elevated CO_2_ resulted in altered expression of genes involved in key functional pathways which were restored to control levels in the intergenerationally treated fish. This “rescue pattern” was predominantly evident in the brain of offspring with sensitive parental phenotype. Specifically, the expression of genes involved in synaptic plasticity and calcium channel activity, initially altered in the developmental treatment, were similar to control levels in the intergenerationally treated fish. Exposure to OA conditions has been shown to affect neural plasticity and neurogenesis in some fish species, which could result in changes in neural circuitry and signaling (Costa et al., 2022; Lai et al., 2017). While neural plasticity could facilitate increased flexibility to environmental changes (Ebbesson & Braithwaite, 2012), it could result in behavioural alterations that have been observed in fish exposed to OA conditions (Schunter et al., 2019). Increased GABAergic signaling is a commonly observed within-generation response to elevated CO_2_ in *A. polyacanthus* (Schunter et al., 2018) and a similar increase in neural signaling pathways was found in the olfactory epithelium of European seabass (*D. labrax*) even after prolonged transgenerational OA exposure for two generations (Cohen-Rengifo et al. 2022). The restoration of OA induced changes in neural signaling processes with previous parental exposure to elevated CO_2_ could indicate elevated intergenerational plasticity of *A. polyacanthus*. In fact, a comparison of six species living in wild CO_2_ seeps with long-term CO_2_ exposures revealed *A. polyacanthus* to have substantially larger brain transcriptional plasticity compared to the other species (Kang et al. 2022). The ’rescue pattern’ of synaptic plasticity genes in intergenerationally treated fish was also accompanied by a similar pattern in the regulatory processes, underlying this plasticity. These regulatory processes are driven by various cell adhesion and cytoskeleton genes that play a role in synaptic plasticity (Gordon-Weeks & Fournier, 2014; Zapara et al., 2000). Therefore, intergenerational CO_2_ exposure potentially restores the dynamic equilibrium of cytoskeletal proteins, thereby re-establishing synaptic signaling processes to control levels.

Additionally, genes involved in transcriptional regulation such as transcription factors and RNA-mediated gene silencers also showed a “rescue pattern” in the brain. Changes in the external environment can trigger reprogramming of transcriptional networks resulting in dynamic regulation of gene expression (Swift & Coruzzi, 2017) as also suggested in wild fish populations naturally exposed to elevated CO_2_ (Petit-Marty et al., 2021). Therefore, these regulatory genes could play a key role in developmental plastic responses to elevated environmental CO_2_ levels, however, these are no longer needed in intergenerationally treated fish revealing acclimation. In the gills, on the other hand, this ‘rescue’ pattern was seen for genes encoding ion transporters in offspring of both behaviorally sensitive and tolerant parents. Notably, potassium channels, known for their significance in pH regulation and association with CO_2_ chemoreception in the gills (Qin et al., 2010), displayed differential expression in the developmental treatment, possibly triggered by hypercapnia. This, in turn, could prompt downstream compensatory responses to maintain pH homeostasis (Suresh et al., 2023, Qin et al., 2010). However, the restoration of expression levels of these genes to control levels in intergenerationally treated fish suggests that intergenerational exposure may enhance and facilitate improved pH regulatory mechanisms.

We also found a new complement of genes to be differentially expressed only when the parents are exposed to the same elevated CO_2_ condition as their offspring, which could further facilitate acclimation of the offspring enabling them to better cope with an elevated CO_2_ environment. Such intergenerational specific transcriptional signature was predominantly observed in the brain and liver of offspring of behaviorally tolerant parents, notably in genes associated with metabolism. Furthermore, genes involved in ion transport leading to bicarbonate retention showed intergeneration-specific differential expression, predominantly in the gills and to a lesser extent in the brain of fish with tolerant parental phenotype. Increasing internal bicarbonate ion concentrations is a commonly observed compensatory response in fish to buffer acid-base disturbance caused by exposure to elevated CO_2_ levels (Heuer & Grosell, 2014). Differential expression of bicarbonate transporters in the intergenerationally treated fish suggests that parental conditioning enables the offspring to more effectively buffer pH changes (Brauner et al., 2019; Chen et al., 2009; Choi, 2012.; Esbaugh, 2017). Therefore, intergenerational CO_2_ exposure could have resulted in transcriptional rearrangements of ion transporters in the offspring, especially in the brain and gills, to maintain pH homeostasis primarily by bicarbonate retention.

Parental conditioning to elevated CO_2_ also increased the offspring’s capacity of energy metabolism indicated by upregulation of genes involved in fatty-acid metabolism in the brain and liver. Additionally, lipid synthesis was also upregulated in the liver. Changes in lipid metabolism has been shown to facilitate acclimation to elevated CO_2_ as well as temperature in previous studies (Liu et al., 2013, Strader et al., 2020, Veilleux et al., 2015). Therefore, upregulation of lipid metabolism in both the brain and liver in intergenerationally treated fish combined with the observed redirection of metabolic carbon fluxes observed when considering the overall CO_2_ response, might indicate a similar switch from carbohydrate to lipid metabolism as the primary metabolic pathway for ATP production in elevated CO_2_ conditions, although further research is needed to confirm this. Furthermore, genes involved in cellular stress response were upregulated in the brain of intergenerationally treated fish with behaviorally tolerant parents. Notably, the cellular stress response pathway was upregulated in the gills and liver in both developmental and intergenerational CO_2_ treatment, but in the brain only in the intergenerational CO_2_ treatment. Given that cellular stress response pathway is an energy demanding process (Strader et al., 2020), the systemic upregulation of this function in all three tissues only in the intergenerationally treated fish could be facilitated by the increased capacity of energy metabolism.

We also found a parental behavioural phenotype to influence the acclimation potential of offspring to elevated CO_2_ conditions. Offspring of tolerant parental phenotype showed a stronger transcriptional response to elevated CO_2_ exposure with higher number of DE genes and larger fold changes across all three tissues. Furthermore, offspring of tolerant parental phenotype showed increased intergenerational plastic responses compared to offspring of behaviorally sensitive parents, with the brain and liver showing the largest signal of intergenerational effects. Parental behavioural phenotype has been shown to influence the offspring brain transcriptional response to elevated CO_2_ in *A. polyacanthus* (Monroe et al., 2021), with behavioural tolerance being heritable (Welch & Munday, 2017). Selection experiments have indicated a genetic basis for individual variation in OA induced responses in a variety of animals (Langer et al. 2009; Parker et al., 2011; Pistevos et al. 2011; Sunday et al. 2011), including the behavioural phenotype to chemical alarm cues in elevated CO_2_ in *A. polyacanthus* (Lehmann et al. 2022). This intraspecific variation in organismal response is key in driving future adaptive evolution. Our results suggest that offspring of parents with a tolerant behavioural phenotype have an increased capacity for intergenerational plasticity and potential acclimation to future ocean acidification conditions.

This study used a systematic approach considering parental behaviour, environment, and multiple tissues to assess molecular responses to future ocean conditions. Gills and liver work together to reduce stress-induced cellular damage in offspring of tolerant parents under all CO_2_ treatments, while in the brain cellular stress response is upregulated only in intergenerationally treated fish. Therefore, parental conditioning to elevated CO_2_ potentially enhances offspring’s stress response at a systemic level. Additionally, prior parental exposure to OA mediates the reprogramming of the offspring transcriptome, impacting energy metabolism in the brain and liver of tolerant-parent offspring and resulting in rescue of altered synaptic signaling pathways, observed in the developmental treatment, in the brain of offspring with behaviourally sensitive parents. Overall, our study reveals how intergenerational plasticity is facilitated from a whole-organism perspective and illustrates how transcriptional changes across multiple tissues integrate to drive the plastic responses of fish to the changing ocean chemistry.

## Author contributions

The experiment was designed and run by MJW and PLM. Molecular lab work was performed by CS and sequenced by TR. SS carried out the transcriptome expression analysis with input from CS. SS lead the writing of the manuscript with input from CS and all authors read, edited and approved the final manuscript.

## Ethics

Sample collection was carried out following all institutional and national law guidelines. The experiment was completed under James Cook University ethics approval A1828.

## Acknowledgements

CS and SS were supported through the HKU start-up to CS. PLM was supported by the ARC Centre of Excellence for Coral Reef Studies and TR was supported by Okinawa Institute of Science and Technology (OIST).

## Data availability

The brain RNA-Seq raw sequences are deposited in NCBI under BioProject ID PRJNA311159. The gills and liver RNA-Seq raw sequences are deposited in NCBI under BioProject ID PRJNA989422 (https://dataview.ncbi.nlm.nih.gov/object/PRJNA989422?reviewer=q3n0q75hbbf2p1v85n0h e3uv86).

