## Supplementary figures for "Cross-talk between tissues is critical for intergenerational acclimation to environmental change"

**Supplemenary Figure 1:**

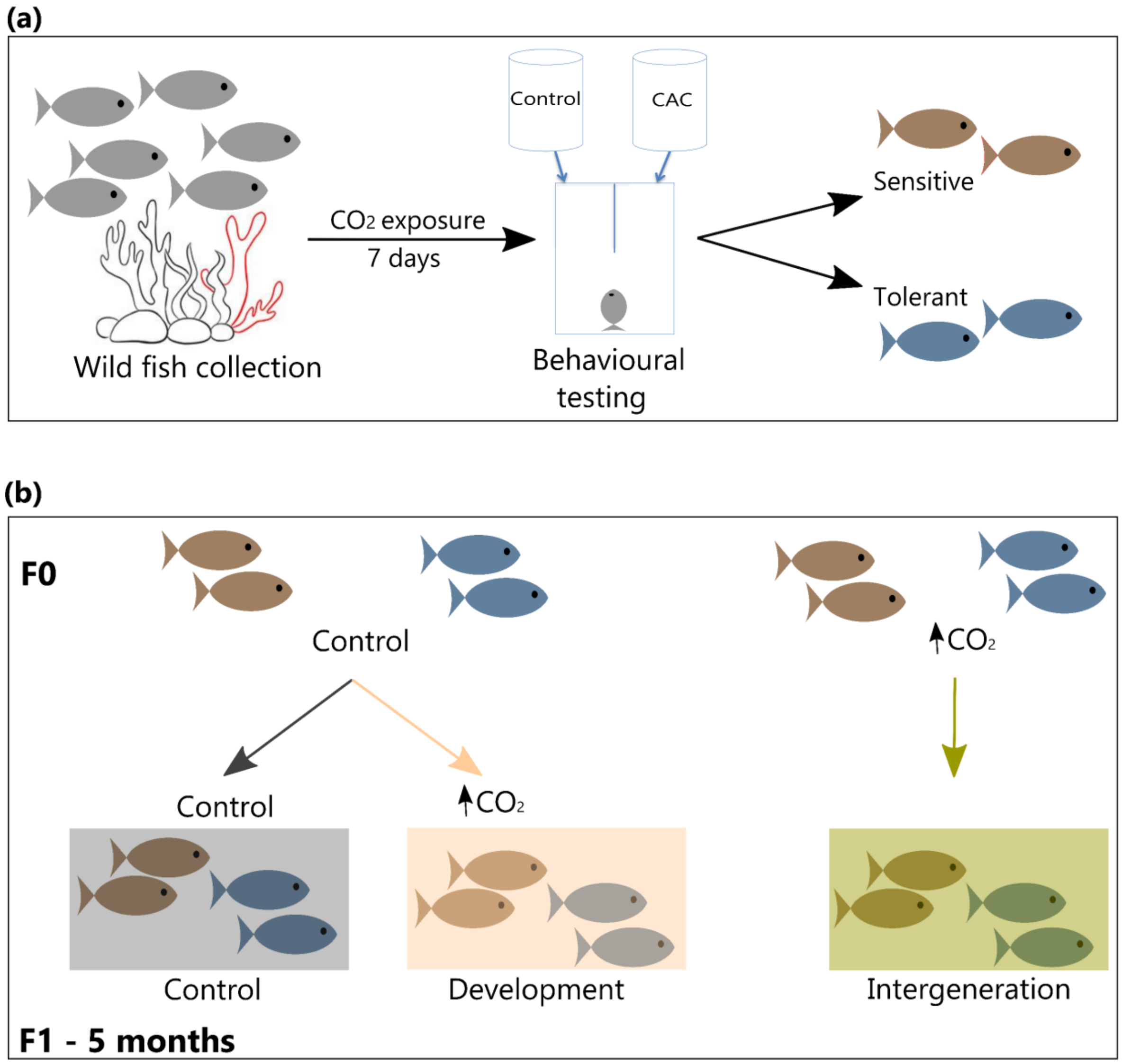


**Figure S1:** Schematic representation of the experimental design including (a) wild fish collection, behavioural testing, (b) creation of breeding pairs, and the $CO_2$ exposure treatments of parents (F0) and offspring (F1). Brown and greyish-blue colours indicates offspring of sensitive and tolerant parents respectively. In figure (b) gray, peach-orange, and olive colours indicate the control, developmental and intergenerational $CO_2$ treatments, respectively.

**Supplemenary Figure 2:**

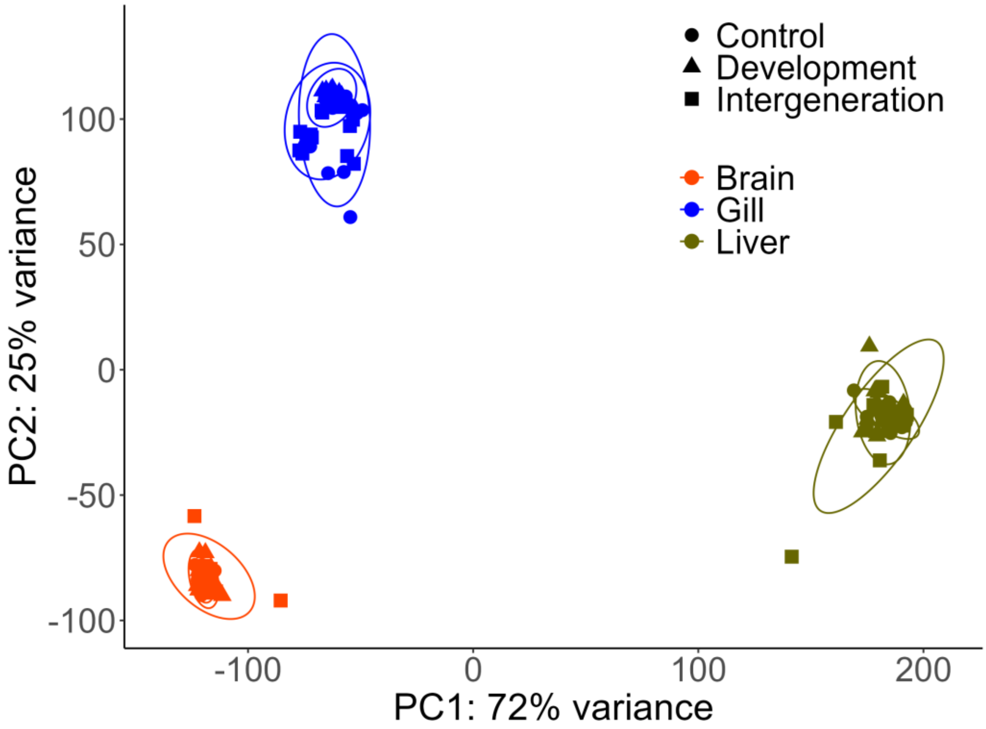


**Figure S2:** Gene expression patterns of offspring of behaviourally sensitive and tolerant parents across all three tissues and treatment conditions. Principal component analysis (PCA) was done using the variance stabilizing transformed (vst) counts off all expressed genes across all the samples (N = 162). Clustering of samples by tissues indicates that the tissues vary greatly in their transcriptional signatures.

**Supplemenary Figure 3:**

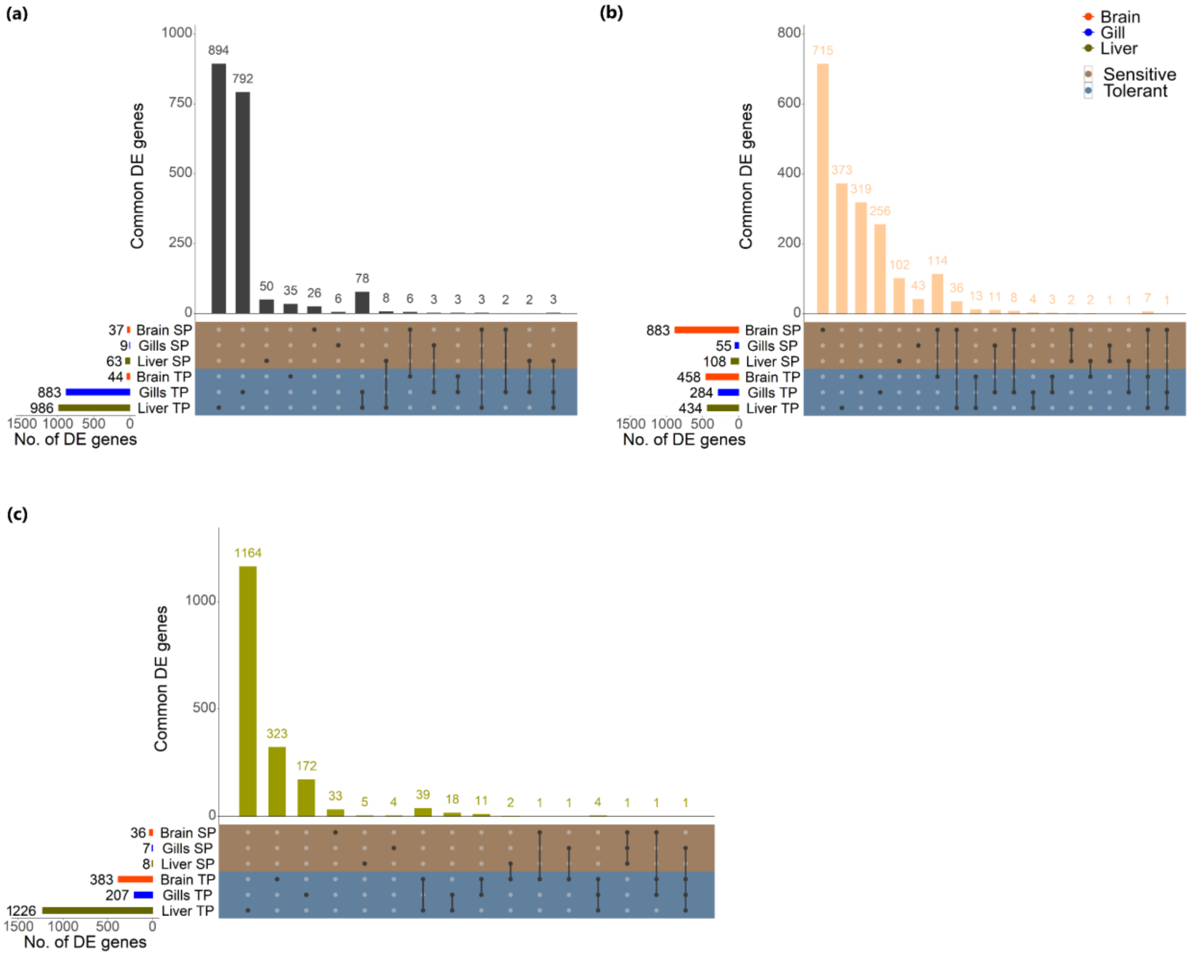


**Figure S3:** upSet plot showing expression paterns of genes involved in (a) overall $CO_2$ resposne, (b) rescue pattern, and (c) intergenration specific resposne across all three tissues. SP indicates samples with a sensitive parental phenotype and TP indicates samples with a tolerant parental phenotype
